# TALE-based C-to-T base editor for multiple homologous genes with flexible precision

**DOI:** 10.1101/2023.02.27.530312

**Authors:** Ayako Hosoda, Issei Nakazato, Hideki Takanashi, Nobuhiro Tsutsumi, Shin-ichi Arimura

## Abstract

Recently a cytidine deaminase-based method for highly efficient C-to-T targeted base editing was developed and has been used with CRISPR-mediated systems. It is a powerful method for genome engineering, although it is prone to off-target effects and has limited targeting scope. Transcription activator-like effector (TALE) -based tools which allow longer recognition sequences than do CRISPR/Cas9 systems, can also be used for targeted C-to-T base editing. Here, we describe a method that efficiently achieved targeted C-to-T substitutions in *Arabidopsis* nuclear genes using cytidine deaminase fused to a TALE DNA-binding domain (TALECD). We used a single pair of TALEs with a novel TALE-repeat unit that can recognize all four DNA bases, especially to allow for variations in the third base of codons in homologous genes. This targeting strategy makes it possible to simultaneously base edit almost identical sites in multiple isoforms of a gene while suppressing off-target substitutions.

## Introduction

Base editing, a genome editing technology, can substitute specific bases for other bases in the target sequence. This tool enables genome modification with minimal changes compared to the nuclease-based gene editing. Base editors, which are composed of a nickase Cas9 (nCas9) and some DNA deaminases, have been used successfully in plants and animals (Komor et al., 2016; Nishida et al., 2016; reviewed in Azameti & Dauda, 2021). However, in some cases, these technologies do not work well, primarily because they have a limited genome-targeting scope due to the requirement for a protospacer adjacent motif (PAM). Undesired bystander base mutations or indel formations can also be obstacles to precise genome editing. Moreover, there is little room to adjust the precision of these methods because they are limited to recognizing around 20-bp-long protospacer sequences, which makes it difficult to prevent off-target mutations.

Recently, a three-component base editor containing a transcription activator-like effector (TALE) DNA-binding domain, a cytidine deaminase domain (CD) of DddA toxin and a uracil glycosylase inhibitor (UGI) was reported to induce C-to-T conversions in mammalian mitochondrial DNA (Mok et al., 2020). DddA is a bacterial toxin which catalyzes the deamination of cytosines within dsDNA. This method uses two TALE proteins, each of which is fused to an inactive half of CD. When the two TALE proteins bind adjacent recognition sequences, the CD is activated and induces the deamination of cytosines in the target window between recognition sequences (Fig. 1a). Organellar (mitochondrial and plastid) genomes are difficult to modify by CRISPR/Cas9 systems because of the difficulty of importing single-guide RNAs (sgRNAs) into organelles (reviewed in Son & Park, 2022). Three groups, including our group, independently developed this TALE-based tool to introduce targeted C-to-T substitutions into the chloroplast and mitochondrial genomes in plants (Kang et al., 2021; Li et al., 2021; Nakazato et al., 2021, 2022). We achieved rapid and efficient base editing, as both mitochondrial and plastid genomes, with 50-100 and more than 1,000 copies per cell, respectively, were homoplasmically edited in one or two generations in *Arabidopsis* (Nakazato et al., 2021, 2022). These successes encouraged us to apply this system to nuclear base editing, designated nTALECD in this study. TALECD works as a dimer, and a pair of DNA-binding domains of TALECD can recognize a broader range (12∼42 bp) in length than can the CRISPR-based system. The longer recognition sequence of a single TALECD pair is expected to result in fewer off-target substitutions than occurs with CRISPR-based sequences.

**Figure 1.**
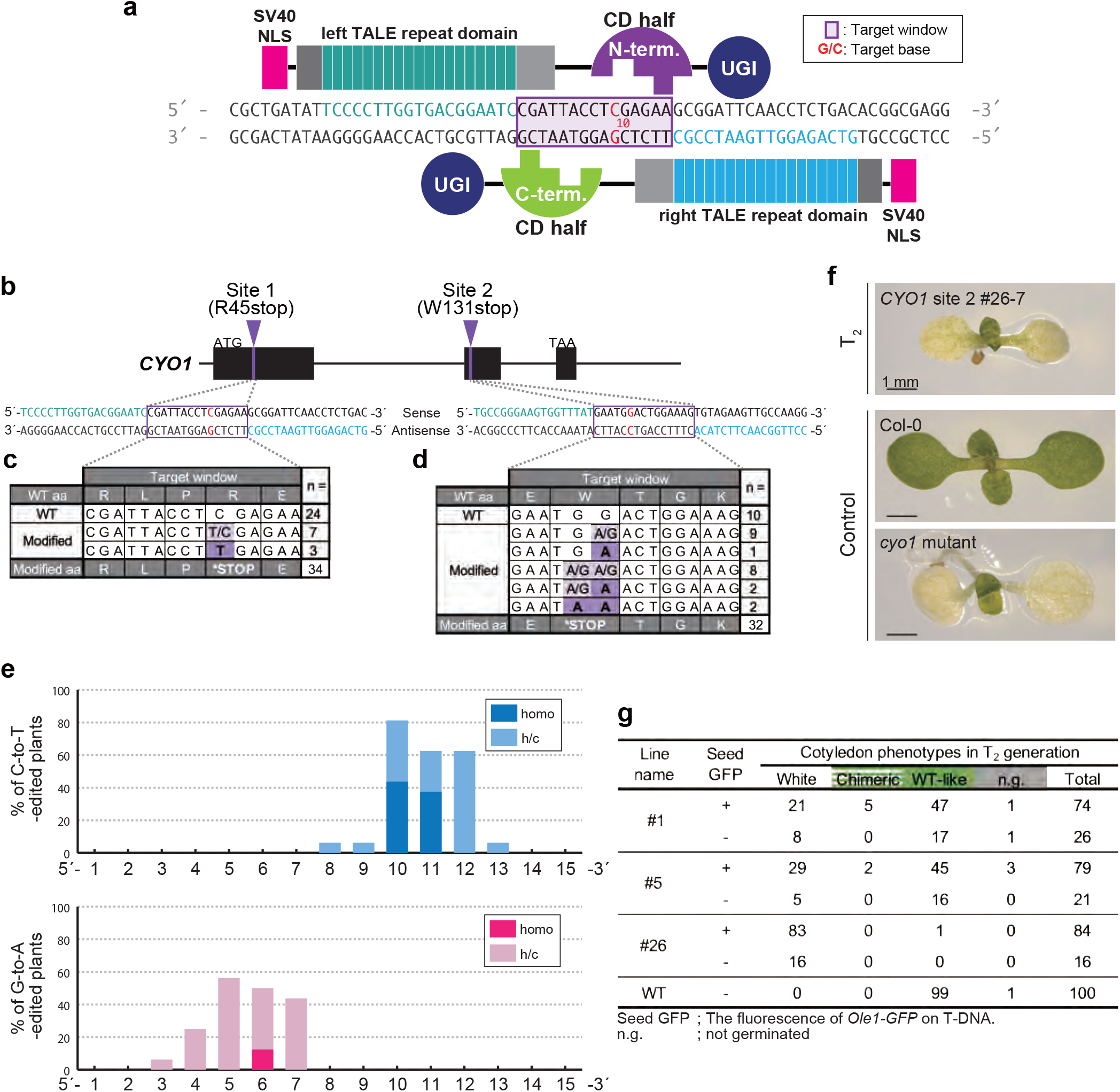
C-to-T base editing in the nuclear genome of *A. thaliana* by nTALECD. (a) Schematic illustration of nTALECD targeting an intended genomic region. Each half of the cytidine deaminase (CD) domain from *Burkholderia cenocepacia* DddA toxin (1,427 aa) split at the 1,397th amino acid, N-half fused to left and C-half to right TALE domain. (b) Schematic representation of the genomic structure of *CYO1*. (c, d) The edited sequences in the target windows and the corresponding numbers of T_1_ plants. *CYO1* site 1 (c) and site 2 (d). (e) Editing activity of 5’
s-tC (upper) or 5’-Ga (lower) contexts at each position in the TC-repeat region. The graph exhibit combined data from four nTALECD (split at the 1,397th amino acid) constructs, shown in Supplemental Fig. 3b-e. Subscripted numbers refer to the positions of cytosines or guanines in the target window counting the DNA nucleotide immediately after the binding site of TALE fused with N-terminal half. h/c; Abbreviation of hetero or chimeric. (f) 10 DAS cotyledon phenotypes. A T_2_ plant, #26-7, has a homoallelic mutation in *CYO1. cyo1* mutant is reported in ref (Shimada et al., 2007). (g) Cotyledon phenotypes of three *CYO1* site 2-targeted T_2_ lines with or without seed GFP fluorescence.

## Results

We constructed plant transformation vectors that express a pair of nTALECDs, each with an SV40 nuclear localization signal (NLS) at the N-terminus (Fig. 1a). In this study, we adopted a 1397N/C nTALECD architecture in which CD is split at the 1,397^th^ amino acid (Mok et al., 2020). Both the N- and C-terminal halves of CD, each fused to DNA-binding domains, were expressed under control of the *RPS5A* promoter (Supplemental Fig. 1) (Tsutsui & Higashiyama, 2016). One of our target genes was *AtCYO1* (*SCO2*, AT3G19220), whose knockout results in albino cotyledons but normally developed true leaves (Shimada et al., 2007). We also targeted two other genes, *AtPKT3* (*PEROXISOMAL 3-KETOACYL-COA THIOLASE 3*, AT2G33150), and *AtMSH1* (*MUTL PROTEIN HOMOLOG 1*, AT3G24320), whose knockouts are not lethal under normal growth conditions (Hayashi et al., 1998; Abdelnoor et al., 2003). For target cytosines, we selected mostly ones at 5’
s-tC sites which DddA toxin strongly prefers (Mok et al., 2020). We also designed nTALECD recognition sites with 14- or 15-bp target windows to place the target cytosine bases in the most probable conversion sites, according to previous reports (Nakazato et al., 2021). Transformation vectors to express pairs of nTALECDs simultaneously were transformed in *Arabidopsis* ecotype Col-0 via an *Agrobacterium*-mediated floral dip (Clough & Bent, 1998).

T_1_ plants at 21 days after stratification (DAS) were subjected to PCR and Sanger sequencing. We observed C:G-to-T:A conversions at the targeted cytosines or complementary guanines, some of which seemed to be homoallelically edited (Fig. 1c, d, Supplemental Table 2, 3). Sanger sequencing revealed the desired nonsense mutations in 10 of 34 transformants in which the *CYO1* site 1 was targeted, and in 22 of 35 transformants in which the *CYO1* site 2 was targeted. Furthermore, while most of the transformants in the first generation had heterozygous mutations, 3 of the 10 site 1-edited transformants and 5 of the 22 site 2-edited transformants had homozygous mutations. C-to-T substitutions were also introduced in the target windows of the other four target sites with similar efficiencies (Supplemental Fig. 2, Supplemental Table 4-7). Interestingly, we detected not only 5’
s-tC conversions but also 5’-tCC-to-tTT conversions. Taken together, these results indicate that TALECD can efficiently catalyze targeted tC-to-tT substitutions in the *Arabidopsis* nuclear genome as well as in organelle genomes.

To investigate which cytosines in a 15-bp target window are prone to be substituted *in vivo* by nTALECD (1397N/C) in the *Arabidopsis* nuclear genome, we newly designed four pairs of nTALECDs targeting a TC-repeat sequence in an intergenic region (Supplemental Fig. 3a). C-to-T substitutions were introduced at specific positions of the target window by each TALECD pair (Supplemental Fig. 3b). The cytosines in the 10th and 11th positions (C10, C11) and the guanine in the 6th position were efficiently substituted to a homoallelic state (Fig. 1g). Furthermore, no indel mutations in the targeted region were detected by Sanger sequencing throughout our study. In contrast, Cas9-mediated base editors can often cause indel byproducts, presumably due to the nicking activity of nCas9 (Choi et al., 2021; Ren et al., 2021). Collectively, these results indicate that nTALECD can efficiently substitute cytosines in a specific region within the target window.

To examine the inheritance of the targeted mutation and the introduced vectors for genome editing, we sowed seeds from three T_1_ individuals that harbored premature stop codons in the target window at *CYO1* site 2 (Supplemental Fig. 4). Among the T_2_ offspring, we observed some mutants with cotyledon-specific albino phenotypes, such as the *cyo1* mutant (Fig. 1f). Although the frequencies of albino cotyledons differed among the three lines, such albino phenotypes were detected in individuals both with and without seed GFP fluorescence (Fig. 1g). This suggests that mutations introduced in *CYO1* were stably inherited in the T_2_ generation, independent of the presence of the nTALECD vectors. Knockout mutant-like phenotypes were also observed in T_2_ plants grown from GFP-negative seeds of the other two target genes (*PKT3* and *MSH1*) (Abdelnoor et al., 2003; Hayashi et al., 1998). These data indicate that nTALECD can efficiently introduce targeted base substitutions in the T_1_ generation and that the mutations were transmissible to the next generation, whether or not the expression cassette of the genome editing enzyme was transmitted.

The DNA-binding domain of TALECD is composed of nearly identical tandem repeats called TALE-repeats, each of which can recognize a single DNA base pair. Each repeat generally has a 102-bp sequence encoding 34 amino acid residues except for the last short repeat. The repeats are identical except for the part encoding the 12th and 13th amino acid residues, called the repeat variable diresidue (RVD), which specifies a particular base (Boch et al., 2009; Moscou & Bogdanove, 2009). For example, the RVDs NI (Asn-Ile) and NG (Asn-Gly) target adenosines and thymines respectively (Fig. 2a). TALECD has an advantage in precision editing since it can recognize up to around 40-bp sequences, whereas Cas9 can recognize sequences of only about half that length. Conversely, the high-specificity of TALECD makes it difficult to mutate homologous genes, which may differ in the target sites. Simultaneous modification of redundant genes is often necessary for trait improvement. A simple option for multiplex gene editing using TALE-based tools is to assemble multiple TALE pairs for each target sequence, but that is laborious and time-consuming. Another way is to target multiple homologous genes with a single TALE pair by using degenerate recognition sequences. To do this, we adopted a new TALE-repeat that recognizes any of the four canonical DNA bases. Yang et al (2014) evaluated the nucleotide recognition preference of 400 possible RVDs, which have all combinations of two amino acids, by *in vitro* assays and identified a significant number of RVDs that target multiple DNA bases. Among those RVDs, RV (Arg-Val) seemed optimal for recognizing all four bases. We thus designate this TALE repeat harboring RV as an “N-recognition repeat”, where N is any base (Fig. 2a).

**Figure 2.**
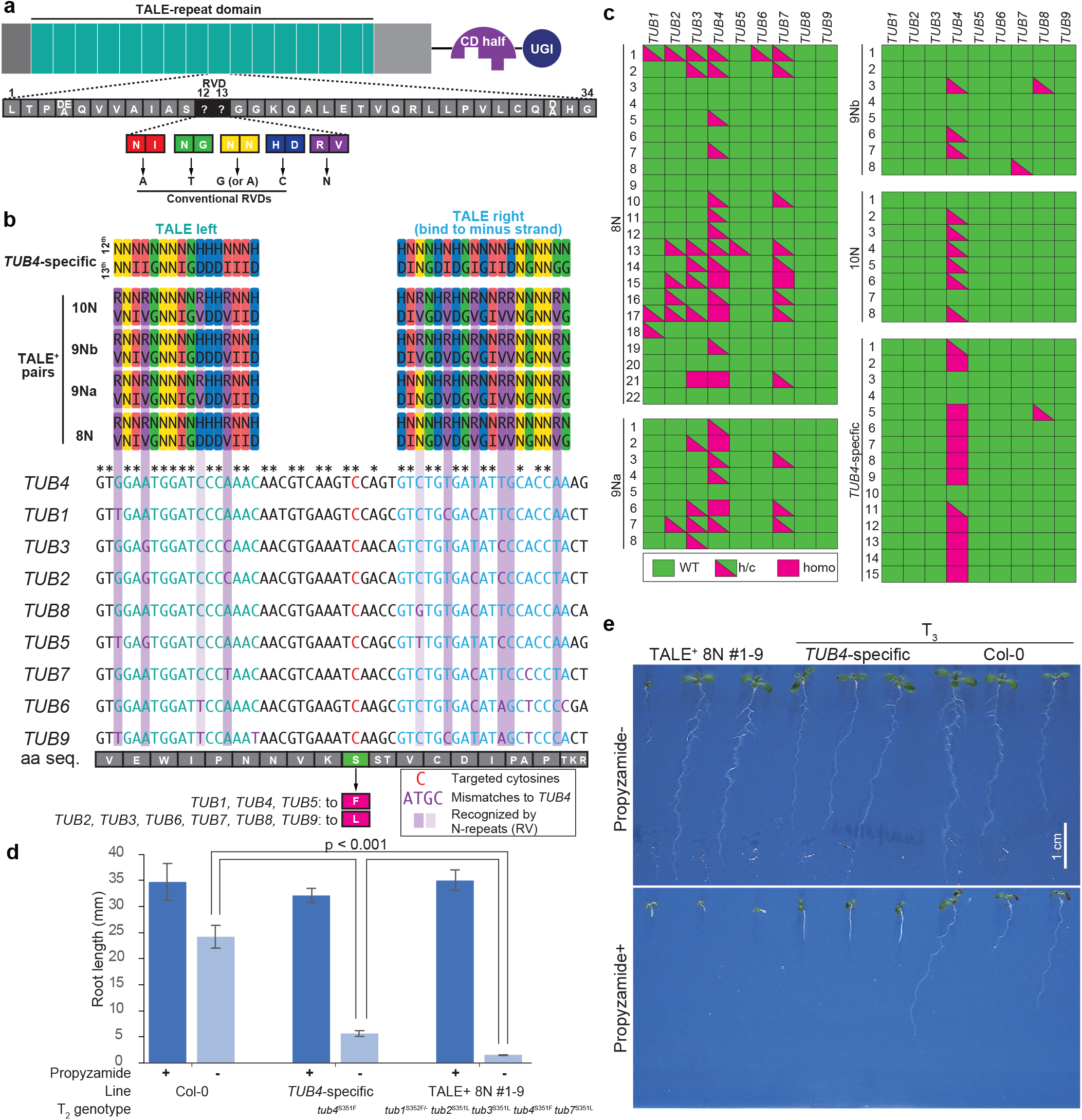
Base editing of β-tubulin genes by a single pair of nTALECD containing N-recognition repeat (TALE^+^). (a) The schematic drawing of the TALE^+^ which possesses TALE-repeats harbouring four canonical RVDs (12th and 13th amino acid residues, NI; Asn-Ile, NG; Asn-Gly, NN;Asn-Asn, HD; His-Asp) which recognize specific bases or new RVD (RV, Arg-Val) acceptable to all four DNA bases (N-recognition RVD). (b) The design of TALE^+^ pairs to target multiple *TUB* genes. The bottom part is an alignment of the target sites of *TUB* genes. The targeted cytosines are shown in red. TALE left- or right-binding sequences are colored in pale green or pale blue, respectively. Base mismatches compared with *TUB4* are colored in purple. The top part shows the structures of the DNA-binding domains of four TALE^+^ pairs, 8N, 9Na, 9Nb and 10N contain 8, 9, 9 and 10 N-repeats in TALE-repeat domain, respectively. The repeat sequences are provided in the RVD code. The purple columns indicate the positions where the target windows differ. (c) Genotypes of targeted cytosines in *TUB4*-specific or TALE^+^-introduced T_1_ plants. (d) Comparison of root length of 7 DAS T_3_ lines with a single mutation only in *TUB4* or mutations in multiple *TUB*s (n ≧10). Each lines were grown on growth medium with or without 3μM propyzamide. (e) 7 DAS seedlings of T_3_ individuals grown on growth medium with or without 3μM propyzamide.

In this study, we attempted multiplex editing of specific serine residues of nine members of the *Arabidopsis* β-tubulin (*TUB*) gene family (Snustad et al., 1992) by single TALECD pairs. Fig. 2b (bottom) shows an alignment of the target sites of nine *TUB* genes. We constructed four TALECD pairs harboring 8, 9, 9, or 10 N-recognition repeats (designated TALE^+^ pairs) (Fig. 2b, top), each targeting cytosines at the 2^nd^ bases of the codons encoding the targeted serine residues. The TALE^+^ pairs are named 8N, 9N, etc., indicating the number of N-recognition repeats in the DNA-binding domains. For example, 8N was designed to target four *TUB* genes (*TUB1, TUB2, TUB3* and *TUB4*) by using eight N-recognition repeats to tolerate eight mismatched nucleotides. The mismatched nucleotides in the alignment are referred to as single nucleotide variants (SNVs). A *TUB4*-specific TALECD pair whose DNA-binding domain is composed of only conventional TALE-repeats was used as a control. Sanger sequencing of all nine *TUB* genes at 14 DAS in the T_1_ generation showed that targeted C-to-T substitutions introduced by *TUB4*-specific TALE pairs were dominantly in *TUB4*, except in one plant that harbored mutations in both *TUB4* and *TUB8* (Fig. 2c, Supplemental Table 8). This result revealed the high specificity of conventional TALE-repeats. All the TALE^+^ pairs except 10N introduced simultaneous C-to-T substitutions in multiple *TUB* genes, whereas 10N modified only *TUB4*. 8N was the most competent in multiple gene targeting, as it could introduce mutations in up to six genes in a T_1_ plant (e.g., 8N #1 in Fig. 2c). Targeted base substitutions introduced by TALE^+^ pairs were predominantly detected as partial ones which are either heteroallelic or chimeric mutations, whereas *TUB4*-specific TALE pairs introduced homoallelic substitutions at *TUB4* in two-thirds of T_1_ plants examined in this study.

To confirm the phenotypes of multiple *TUB* mutants, we grew T_3_ plants originating from a T_1_ plant expressing 8N with the targeted mutations in *TUB1, TUB2, TUB3, TUB4* and *TUB7*. In the T_2_ generation, the mutations were homozygous for all loci except those in *TUB1*. The phenotypes of the multiple *TUB-*mutated T_3_ plants were compared to a *TUB4* single mutant originating from a *TUB4*-specific transgenic T_1_ plant. Seed GFP fluorescence was not observed in the T_3_ plants, and was presumably removed by the T_2_ generation. A point mutation in TUB4 resulting in an S-to-F substitution at residue 351 is known to raise sensitivity to propyzamide, a microtubule-depolymerizing agent (Ishida et al., 2007). When the mutants and wild-type plants were grown on normal medium, their root lengths at 7 DAS were not significantly different (Fig. 2d, e). However, in the presence of 3 μM propyzamide, root elongation in each of two *tub* mutant lines examined was more inhibited than it was in the wild-type. Root elongation of the *tub4* single mutants was moderately inhibited, whereas it was strongly inhibited in *tub* multiple mutants. By allowing SNVs in the alignments of target genes by N-recognition repeats, we could successfully introduce the desired amino acid mutations at homologous positions in multiple *TUB* genes in the T_1_ generation using a single TALE^+^ pair. These mutations were stabilized by the T_3_ generation.

## Discussion

The preceding results demonstrate that TALECD can be used to modify a plant nuclear genome. When transformed with TALECDs composed only of conventional TALE-repeats, T_1_ individuals which harbored homoallelic C:G > T:A substitutions at tC sites were obtained for all target sites. Although these results were mostly limited to tC sites, the base editing efficiency in *Arabidopsis* seemed comparable to that of two CRISPR/Cas9 based cytosine base editors (CBEs) with rAPOBEC1 or PmCDA1 (Choi et al., 2021). Mok et al. recently developed an enhanced DddA toxin that could edit mitochondrial DNA not only at tC sites but also at non-tC sites (Mok et al., 2022). In addition, the targeting scope of TALECD is thought to be broad because of its flexibility of target design and variability of the length of the recognition sequence. Taken together, these findings show that TALECD has the potential to be a versatile tool for nuclear genome editing.

In this study, we took advantage of the N-recognition repeat harboring RV (Arg-Val) as the repeat variable diresidue (RVD), which recognizes four DNA bases and was used to broaden the recognition sequence for multiplex gene editing by the TALE system. Multiplex genome editing is usually achieved by simultaneously using different single-guide RNAs (sgRNAs) in the CRISPR system. However, this increases the risk of off-target effects. By contrast, multiplex editing using a single TALEN or TALECD pair with N-recognition repeats can efficiently edit on-target sites, suppressing off-target modifications by using longer (up to 40-bp) recognition sequences. In the *TUB4*-specific TALE pair, for example, a recognition sequence length of 35 bp (requiring 35 TALE repeats) could reduce the expected frequency of the specific target sequence, by about six orders of magnitude compared to CRISPR/Cas9 which recognizes 20-bp sequences. This calculation was based on an assumption that NN (Asn-Asn) has a high preference for both guanine and adenine. Also, even in the case of the TALE^+^ 8N, whose specificity relies on a total of 27 (=35-8) conventional TALE-repeats, the off-target frequency can be two orders of magnitude less than those of CRISPR/Cas9. This indicates that multiplex genome editing can be achieved by the TALE system with high precision. On the other hand, in our experiments, multiple mutations were less common in 9Na, 9Nb, and 10N than those in 8N, suggesting that the ratio of N-recognition repeats to total TALE-repeats needs to be adjusted to each case.

The TALE system might also be able to edit nuclear genomes without using DNA vectors or RNA molecules. In contrast, RNA-free genome editing is almost impossible with the CRISPR/Cas system, which normally requires RNAs for genome targeting. Plants modified by the direct injection of TALECD proteins should not be regarded as GMO plants (and thus not be subject to GMO regulations) in some countries like Japan, because they do not contain any exogenous nucleic acid. Developing nTALECD and other genome editing technologies with precisions and other characteristics that differ from those of CRISPR-based technologies will expand the possibilities of targeted base substitutions, which, in turn, will have many applications in plant science and breeding.

## Materials and Methods

### Construction of Ti plasmids

To assemble the DNA binding domains of nTALECD, we performed two-step cloning using a Platinum Gate TALEN kit (Addgene, #1000000043). In the assembly-step 1, we performed the ligation of fragments from module plasmids in the presence of BsaI-HF (New England BioLabs, Ipswich, MA, USA) to generate array plasmids, each of which consisted of up to four TALE-repeats. Next, array plasmids were cloned into entry vectors which had the coding sequences of the C-terminal repeat and either N-terminus or C-terminus domain of cytidine deaminase fused to uracil glycosylase inhibitor. After assembly-step 2, the entry vectors harboring the whole coding sequences of left or right TALECD, the entry vector 2 and the destination vector were assembled by a multisite Gateway LR reaction (ThermoFisher Scientific, Waltham, MA, USA). The end product of this step is a Ti plasmid tandemly expressing left and right TALECDs with an NLS at their N-terminus. Other details are described in Vector construction. Primers are listed in Supplemental Table 1.

### Target design of TALECD

The recognition sequences were designed across the target window. The 5’
s end of recognition sequences was set adjacent to thymine because the DNA-binding domains of Platinum Gate TALEN require the thymine to bind the target strongly (Sakuma et al., 2013).

RV (12th amino acid residue; arginine, 13th; valine) was chosen for the N-recognition repeat among a total of 400 types of potential RVDs screened in Yang et al., (2014). The details of the cloning of module plasmids of RV are in Vector Construction. In the TALE^+^ study, nine *TUB* genes expressed in *A. thaliana* Col-0 were targeted. Those genes exhibit 89-96% protein sequence identity but with a significant number of single nucleotide variants (SNVs), including synonymous substitutions in their coding sequences. Desired point mutation, *tub4*^S351F^ was known to locate at the interface of tubulin heterodimers and the serine residue is conserved in all *TUB* genes. C-to-T substitutions of the second base of the codon introduce the following amino acid substitutions: *TUB1, TUB4, TUB5*; Ser > Phe, *TUB2, TUB3, TUB6, TUB7, TUB8, TUB9*; Ser > Leu.

### Transformation and Screening of Transformants

All Ti plasmids were introduced to *Agrobacterium tumefaciens* strain C58C1 (pMP90) and transformed to *A. thaliana* ecotype Col-0 by floral dip. Since each Ti plasmid has a marker gene cassette of Ole1 pro::Ole1-GFP (Shimada et al., 2010) expressed specifically in seeds, transgenic T_1_ seeds were selected by observing seed GFP fluorescence. After Sterilized in 70% (v/v) Ethanol and 5% (v/v) sodium hypochlorite, seeds were sown on the half-strength MS medium containing 125 μg/mL of claforan.

### Growth Conditions and Genotyping

After stratification at 4°C, seeds were cultured in a growth chamber at 22°C under long-day conditions (16 h light, 8 h dark). An emerging true leaf of each T_1_ plant was harvested at 14 DAS or 21 DAS, and subjected to total DNA extraction by boiling in rapid extraction buffer containing 100mM Tris (pH9.5) and10mM EDTA (pH8.0). The genomic region including each target window was amplified with target-specific primers (Supplemental Table 1). The purified PCR products were directly sequenced by Eurofins Genomics (city, Japan).

T_2_ plants were sown on half-strength MS medium without claforan and grown under the same condition as above.

### Vector construction

The Gateway destination vector harboring *Arabidopsis* RPS5A promoter, SV40NLS, and HSP terminator (Addgene #193370) was prepared using In-Fusion Cloning system (Clontech, Mountain View, CA, USA). Two DNA fragments were independently amplified with two primer sets (NLS_Fw1/Rv1, NLS_Fw2/Rv2) using pDest_pK7WG2_pRPS5A_CTP OleGFP (Addgene #171723) as a PCR template. The amplified fragments were cloned into linearized pDest_pK7WG2_pRPS5A_CTP OleGFP digested by AatⅡ and PacI (New England BioLabs).

The Gateway attR4-attR3 entry vector harboring the *RPS5A* promoter, SV40NLS, HSP terminator (Addgene #193371) were generated by site-directed mutagenesis. The full-length fragment of E2_pENTR_R4R3_T CTP (Addgene #171736) were amplified using primers with SV40NLS sequence added (NLS_E2_Fw/Rv). The PCR product was incubated with DpnI (NEB, USA) in 1 hour at 37°C to digest the template plasmid.

TALE-repeats for N-recognition use RV as the RVD. To construct p1RV (Addgene, #193372), a Platinum Gate TALEN assembly vector with this RVD, two DNA fragments were independently amplified with primer sets (RV_Fw1/Rv1, RV_Fw2/Rv2) and p1HD (Addgene #50664) as a PCR template. These fragments were cloned into linearized p1HD digested by PvuII-HF (NEB) using an In-Fusion Cloning system. p2RV (Addgene #193373), p3RV (Addgene #193374) and p4RV (Addgene #193375) were constructed in the same way based on p2HD (Addgene #50668), p3HD (Addgene #50672) and p4HD (Addgene #50676), respectively.

### Image processing

Plant images were taken by Leica MC 170 HD (Leica, Germany), OLYMPUS OM-D E-M5 (Olympus, Japan) and ChemiDoc MP Imaging system (BIO-RAD, USA). These images were processed with Adobe Photoshop 2021. All figures and tables were organized with Adobe Illustrator 2021, Adobe Photoshop 2021 and Microsoft Excel for Mac.

## Acknowledgements

We thank Yoshiko Tamura and Reiko Masuda for their help in the experiments.

## References

Abdelnoor RV, Yule R, Elo A, Christensen AC, Meyer-Gauen G, & Mackenzie SA (2003) Substoichiometric shifting in the plant mitochondrial genome is influenced by a gene homologous to MutS. Proc Natl Acad Sci. 100: 5968–5973

Azameti MK, Dauda WP (2021) Base editing in plants: applications, challenges, and future prospects. Front Plant Sci. 12: 664997

Boch J, Scholze H, Schornack S, Landgraf A, Hahn S, Kay S, Lahaye T, Nickstadt A, Bonas U (2009) Breaking the Code of DNA Binding Specificity of TAL-Type III Effectors. Science 326: 1509–1512

Choi M, Yun JY, Kim JH, Kim JS, Kim ST (2021) The efficacy of CRISPR-mediated cytosine base editing with the RPS5a promoter in Arabidopsis thaliana. Sci Rep. 11: 8087

Clough SJ, Bent AF (1998) Floral dip: A simplified method for Agrobacterium-mediated transformation of Arabidopsis thaliana. Plant J. 16: 735–743

Hayashi M, Toriyama K, Kondo M, Nishimura M (1998) 2,4-Dichlorophenoxybutyric acid– resistant mutants of Arabidopsis have defects in glyoxysomal fatty acid β-oxidation. Plant Cell 10: 183–195

Ishida T, Kaneko Y, Iwano M, Hashimoto T (2007) Helical microtubule arrays in a collection of twisting tubulin mutants of Arabidopsis thaliana. Proc Natl Acad Sci. 104: 8544– 8549

Kang BC, Bae SJ, Lee S, Lee JS, Kim A, Lee H, Baek G, Seo H, Kim J, Kim JS (2021) Chloroplast and mitochondrial DNA editing in plants. Nat Plants 7: 899–905

Komor AC, Kim YB, Packer MS, Zuris JA, Liu DR (2016) Programmable editing of a target base in genomic DNA without double-stranded DNA cleavage. Nature 533: 420–424

Li R, Char SN, Liu B, Liu H, Li X, Yang B (2021) High-efficiency plastome base editing in rice with TAL cytosine deaminase. Mol Plant 14: 1412–1414.

Mok BY, de Moraes MH, Zeng J, Bosch DE, Kotrys AV, Raguram A, Hsu F, Radey MC, Peterson SB, Mootha VK, Mougous JD, Liu DR (2020) A bacterial cytidine deaminase toxin enables CRISPR-free mitochondrial base editing. Nature 583: 631– 637

Mok BY, Kotrys AV, Raguram A, Huang TP, Mootha VK, Liu DR (2022) CRISPR-free base editors with enhanced activity and expanded targeting scope in mitochondrial and nuclear DNA. Nat Biotechnol. 40: 1378–1387

Moscou MJ, Bogdanove AJ (2009) A simple cipher governs DNA recognition by TAL effectors. Science 326: 1501

Nakazato I, Okuno M, Yamamoto H, Tamura Y, Itoh T, Shikanai T, Takanashi H, Tsutsumi N, Arimura S (2021) Targeted base editing in the plastid genome of Arabidopsis thaliana. Nat Plants 7: 906–913

Nakazato I, Okuno M, Zhou C, Itoh T, Tsutsumi N, Takenaka M, Arimura, S (2022) Targeted base editing in the mitochondrial genome of Arabidopsis thaliana. Proc Natl Acad Sci. 119: e2121177119

Nishida K, Arazoe T, Yachie N, Banno S, Kakimoto M, Tabata M, Mochizuki M, Miyabe A, Araki M, Hara KY, Shimatani Z, Kondo A (2016) Targeted nucleotide editing using hybrid prokaryotic and vertebrate adaptive immune systems. Science 353: aaf8729

Ren Q, Sretenovic S, Liu G, Zhong Z, Wang J, Huang L, Tang X, Guo Y, Liu L, Wu Y, et al. (2021) Improved plant cytosine base editors with high editing activity, purity, and specificity. Plant Biotechnol J. 19: 2052–2068

Sakuma T, Ochiai H, Kaneko T, Mashimo T, Tokumasu D, Sakane Y, Suzuki K, Miyamoto T, Sakamoto N, Matsuura S, Yamamoto T (2013) Repeating pattern of non-RVD variations in DNA-binding modules enhances TALEN activity. Sci Rep. 3: 3379

Shimada H, Mochizuki M, Ogura K, Froehlich JE, Osteryoung KW, Shirano Y, Shibata D, Masuda S, Mori K, Takamiya K (2007) Arabidopsis cotyledon-specific chloroplast biogenesis factor CYO1 is a protein disulfide isomerase. Plant Cell 19: 3157–3169

Shimada TL, Shimada T, Hara-Nishimura I (2010) A rapid and non-destructive screenable marker, FAST, for identifying transformed seeds of Arabidopsis thaliana. Plant J. 61: 519–528

Snustad DP, Haas NA, Kopczak SD, Silflow CD (1992) The small genome of Arabidopsis contains at least nine expressed beta-tubulin genes. Plant Cell, 4: 549–556

Son S, Park SR (2022) Challenges facing CRISPR/Cas9-based genome editing in plants. Front Plant Sci. 13: 902413

Tsutsui H, Higashiyama T (2016) pKAMA-ITACHI vectors for highly efficient CRISPR/Cas9-mediated gene knockout in Arabidopsis thaliana. Plant Cell Physiol. 58: 46–56

Yang J, Zhang Y, Yuan P, Zhou Y, Cai C, Ren Q, Wen D, Chu C, Qi H, Wei W (2014) Complete decoding of TAL effectors for DNA recognition. Cell Res. 24: 628–631

